# The *Clostridioides difficile* S-Layer Protein A (SlpA) serves as a general phage receptor

**DOI:** 10.1101/2022.09.19.508581

**Authors:** Alexia L.M. Royer, Andrew A. Umansky, Marie-Maude Allen, Julian R. Garneau, Maicol Ospina-Bedoya, Joseph A. Kirk, Gregory Govoni, Robert P. Fagan, Olga Soutourina, Louis-Charles Fortier

**Affiliations:** Department of Microbiology and Infectious Diseases, Faculty of Medicine and Health Sciences, Université de Sherbrooke, Sherbrooke, Québec, Canada; The Florey Institute, School of Biosciences, University of Sheffield, UK; AvidBiotics, South San Francisco, California, USA; Université Paris-Saclay, CEA, CNRS, Institute for Integrative Biology of the Cell (I2BC), Gif-sur-Yvette, France

## Abstract

Therapeutic bacteriophages (phages) are being considered as alternatives in the fight against *Clostridioides difficile* infections. To be efficient, phages should have a wide host range, but the lack of knowledge about the cell receptor used by *C. difficile* phages hampers the rational design of phage cocktails. Recent reports suggested that the *C. difficile* surface layer protein A (SlpA) is an important phage receptor, but clear and unambiguous experimental evidence is lacking. Here, using the epidemic R20291 strain and its FM2.5 mutant derivative lacking a functional S-layer, we show that the absence of SlpA renders cells completely resistant to infection by phiCD38-2, phiCDIII and phiCD146, which normally infect the parental strain. Complementation assays with 12 different Slayer Cassette Types (SLCTs) expressed from a plasmid revealed that SLCT-6 also allowed infection by phiCD111, and SLCT-11 enabled infection by phiCD38-2 and phiCD146. Of note, expression of SLCTs 1, 6, 8, 9, 10 or 12 conferred susceptibility to infection by 5 myophages that normally do not infect the R20291 strain, namely phiMMP02, phiMMP03, phiMMP04, phiCD506 and phiCD508. Adsorption assays showed that >50% adsorption was required for productive phage infection. Altogether, our data suggest that many phages use SlpA as their receptor and most importantly, morphologically distinct phages of the *Siphoviridae* and *Myoviridae* families target SlpA despite major differences in their tail structures. Our study therefore represents an important breakthrough in our understanding of the molecular interaction between *C. difficile* and its phages.

**IMPORTANCE:** Phage therapy represents an interesting alternative to treat *Clostridioides difficile* infections because contrary to antibiotics, most phages are highly species-specific, thereby sparing the beneficial gut microbes that protect from infection. However, currently available phages against *C. difficile* have a narrow host range and target members from only one or a few PCR ribotypes. Without a clear comprehension of the factors that define host specificity, and in particular the host receptor recognized by phages, it is hard to develop therapeutic cocktails in a rational manner. In our study, we provide clear and unambiguous experimental evidence that SlpA is a common receptor used by many siphophages and myophages. Although work is still needed to define how a particular phage RBP binds to a specific SLCT, identification of SlpA as a common receptor is a major keystone that will facilitate the rational design of therapeutic phage cocktails against clinically important strains.

## INTRODUCTION

With the increasing antibiotic resistance worldwide, there is a regained interest in phage therapy nowadays (1). Bacteriophages (or phages) have the advantage of being highly specific toward their bacterial host, thereby sparing surrounding bacterial species. Since broad spectrum antibiotics often lead to collateral damages, more targeted therapeutics are needed, especially in the context of gastrointestinal infections (2). *Clostridioides difficile* is one of the Gram-positive pathogens for which phage therapy has been proposed as a potential therapeutic alternative (3, 4). *C. difficile* is the main cause of antibiotic-associated diarrhea in hospitals and therapeutic solutions are limited, particularly in the context of recurrent infections (2). *C. difficile* takes advantage of the microbiota dysbiosis caused by broad-spectrum antibiotics to colonize and persist in the gastrointestinal tract. This opportunistic bacterium induces severe and often recurrent intestinal infections driven by the production of toxins (TcdA and TcdB). Since the emergence of the ribotype 027 epidemic strain, *C. difficile* is considered as an urgent threat by the US Centers for Disease Control and Prevention (Report on antibiotic resistance, 2019) (5).

*C. difficile* is susceptible to infection by two structurally different types of phages from the order *Caudovirales*, the *Myoviridae* and *Siphoviridae*. The former is composed of phages with non-flexible and contractile tails whereas the later comprises phages with long, flexible, and non-contractile tails. Like many other phages, phages infecting *C. difficile* generally have a narrow host range (3, 6, 7), which can be the result of multiple factors including the presence of an endogenous Clustered Regularly Interspaced Short Palindromic Repeats (CRISPR) (8), restriction-modification systems (9), phage superinfection exclusion mechanisms such as CwpV (10) and repressor-mediated resistance provided by endogenous prophages (11, 12). Another reason that can explain a narrow host range is the absence of a suitable host receptor at the surface of the target bacteria. The study of phage receptors is therefore crucial to better select suitable therapeutic phages, and because phage resistance often comes from mutation of the receptor (12).

Recognition and binding to a specific receptor at the surface of a bacterial cell is the first and key step for a successful bacteriophage infection. The adsorption process involves close interaction between a cell surface component and a phage counterpart, generally located at the tip of the tail in the case of tailed phages. This adsorption generally occurs in two-steps: the first one involves reversible binding of the phage to the host cell surface through tail fibers or other decorations, followed by the irreversible binding of the phage receptor-binding protein (RBP) to the same host receptor, a different one or both (6, 13).

Substantial structural work has been done on model phages like the *Myoviridae* phages T4 (14) and A511 (15), as well as the *Siphoviridae* phages λ (16), SPP1 (17) and p2 (18, 19). This allowed the identification of key tail components involved in receptor recognition, including baseplate components, tail fibers and the RBP, which plays a central role in receptor recognition (20). Because phage genomes are highly modular and genomic synteny is generally observed, it is now possible to predict with some confidence the RBP and other tail components that compose the host-recognition machinery by homology searches in public repositories. The identification of phage host receptors is more complex, however. Isolation of a spontaneous bacteriophage-insensitive mutant (BIM) is one of the best ways to identify such receptors (21, 22). A large diversity of phage receptors has been characterized to date, revealing bacterial evolutionary strategies to overcome phage infection. Such receptors include proteins, polysaccharides, lipopolysaccharides, capsules, pili, and flagella (23). Much work was done in Gram-negative bacteria, and less is known about phage receptors in Gram-positive bacteria (20). Due to the constant arms races between phages and bacteria, there is a large variability among phage receptors and RBPs, and as such, phage-host interactions remain poorly understood for most bacterial species (24, 25). In *C. difficile*, the identity of phage receptors was unknown until recently, and current data point to the cell-surface protein SlpA as being an important phage receptor.

The *C. difficile* cell surface is composed of a dense proteinaceous array, called the surface layer or “S-layer”, which contains various cell wall associated proteins among which SlpA is the most abundant (26). The S-layer was shown to play various roles in cell adhesion and pathogenesis, immunity, permeability of the bacterial cell and motility (27–29). The SlpA protein is post-translationally cleaved into two fragments by the cell wall associated protease Cwp84 and secreted at the cell surface (26). A high molecular weight (HMW) moiety is attached to the cell wall through cell wall binding domains while a second low molecular weight (LMW) fragment is reassociated to the HMW (30). To date, 14 different SlpA isoforms (also named SLCTs for S-Layer Cassette Types), have been described where the HMW part of the protein is the most conserved and the LMW portion, which is the most exposed at the bacterial surface, is the most variable (31, 32).

Diffocins, are phage tail-like R-type bacteriocins structurally resembling *Myoviridae* phage tails (33). A recent study revealed that diffocins engineered with a Myovirus-derived RBP, called Avidocin-CD291.2, attach to and lyse *C. difficile* cells upon binding to SlpA, which acts as their main receptor (34). While studying Avidocin-CD291.2, complete resistance was observed in two spontaneous mutants of the *C. difficile* epidemic strain R20291 (ribotype 027). These mutant strains, FM2.5 and FM2.6, carry point mutations introducing either a premature stop codon or causing a translational frameshift leading to severe truncation of SlpA. Complementation of the FM2.5 mutant with different SLCTs carried on plasmids rescued susceptibility to killing by different Avidocin-CD (34). Genetic engineering of these Avidocin-CD by replacement of their RBP with a predicted prophage RBP conferred the prophage’s host range to the Avidocin-CD, which led the authors to suggest that SlpA is likely the receptor used by phages as well (34, 35). Another study, based on gel retardation assays with S-layer extracts, also suggested that the S-layer was targeted by the *C. difficile* phage φHN10 (36). Whittle *et al* showed that introduction into strain 630 of plasmids carrying either SLCT-6 or SLCT-H2/6 allowed adsorption of phage φCD1801, a *Myoviridae* phage targeting ribotype 078 isolates but that does not infect strain 630 (ribotype 012) (37). Finally, another recent report suggests that phiCDHS-1, a siphophage infecting the R20291 strain, also recognizes SlpA because it could not infect the FM2.5 mutant (38). Altogether, these data are supporting the idea that SlpA is a receptor for certain phages, but in all the above studies, the evidence available is either indirect or incomplete.

In this study, we sought to fill this knowledge gap by providing unambiguous experimental demonstration that SlpA is a general phage receptor in *C. difficile*. To achieve this, we used the R20291 FM2.5 *slpA* mutant strain complemented with 12 different SlpA isoforms and a collection of 8 different phages. Productive infections and adsorption assays allowed us to clearly demonstrate that SlpA is a general receptor used by many phages to infect *C. difficile*. The data described herein provide a solid foundation for future work aiming at better characterizing phage-host interactions in *C. difficile*.

## RESULTS

### Loss of SlpA confers resistance to phage infection

Recent studies pointed toward a role of the *C. difficile* surface layer protein SlpA in phage infection (34, 36, 39). However, only indirect evidence based on engineered Avidocin-CD (34) or incomplete evidence based only on adsorption assays (36, 39) are currently available. Furthermore, previous studies suggesting that the S-layer or SlpA is a phage receptor were conducted with *Myoviridae* phages only. Hence, whether SlpA also serves as a receptor for *Siphoviridae* phages remains to be demonstrated. This question is of great importance since the tail architecture of *Myoviridae* and *Siphoviridae* phages is very different and there is no indication that both phage families use the same type of receptor to infect their host.

We have previously shown that the wildtype (WT) R20291 epidemic strain is susceptible to infection by the *Siphoviridae* phages phiCD38-2, phiCD111 and phiCD146 (7). To determine whether SlpA could also serve as a receptor for *Siphoviridae* phages, we tested the susceptibility of the FM2.5 *slpA* mutant strain described by Kirk *et al* (34) to infection by these three well-described phages.

As a first step, glycine extracts prepared from the WT R20291 and the FM2.5 *slpA* mutant confirmed the absence of SlpA from the cell surface of the latter (Fig. 1). Then, using a spot test assay, we challenged the FM2.5 mutant with the three siphophages and used the WT R20291 strain as a control (Fig. 2, left panel). The absence of SlpA from the cell surface led to complete resistance to the three phages and no phage mutants could be observed in all our assays (Fig. 2, center panel). To confirm that SlpA is the receptor used by *Siphoviridae* phages, we performed complementation assays. Introduction of a plasmid carrying a gene encoding the SLCT-4 in the R20291 background (that naturally expresses SLCT-4) was previously shown to lead to homologous recombination with the chromosomal copy. Hence, we used the revertant strain FM2.5RW, in which the chromosomal mutated gene has been replaced with a “watermarked” functional *slpA* gene containing two synonymous mutations (34). Expression of SlpA in the FM2.5RW strain was confirmed by glycine extraction of cell surface proteins (Fig. 1, 4^th^ lane), and spot test assays confirmed full restoration of susceptibility to phage infection (Fig. 2, right panel). Together, these results clearly suggest that SlpA is also a receptor for these *Siphoviridae* phages.

**FIGURE 1.**
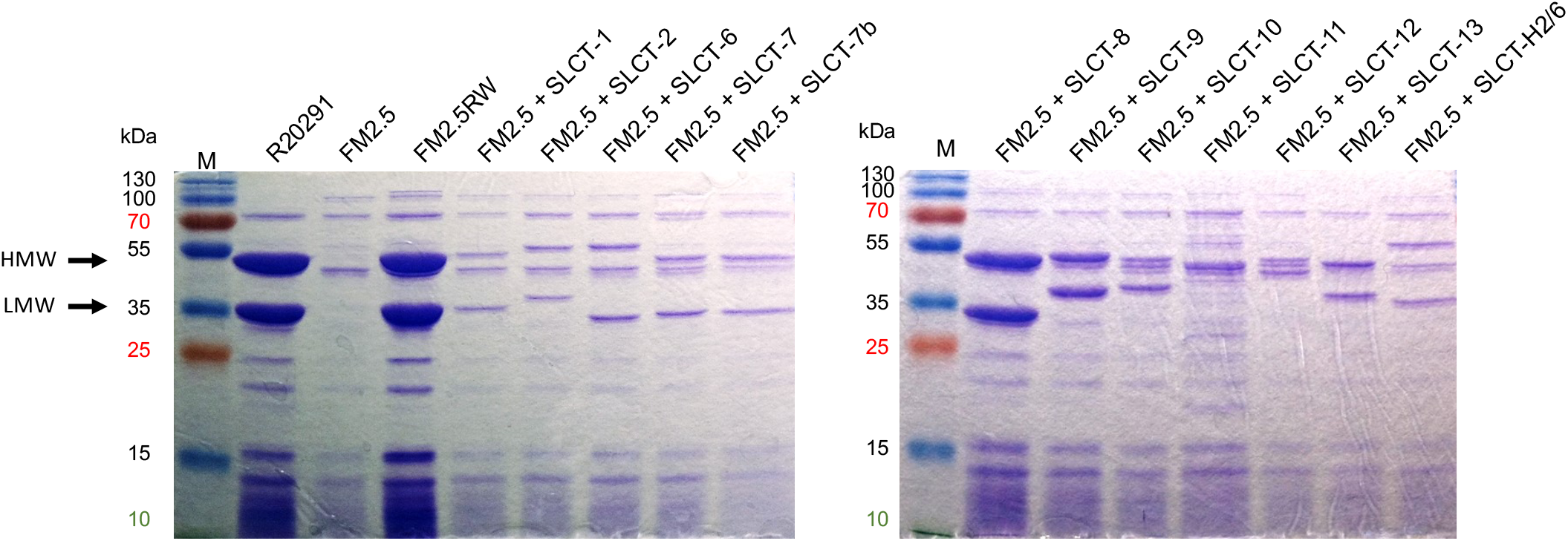
Coomassie-stained 12% SDS-PAGE showing glycine extractions of surface proteins from WT R20291, FM2.5 *slpA* mutant, FM2.5RW watermarked revertant and the FM2.5 mutant complemented with a plasmid encoding one of the 12 other SLCTs tested. Expression of the SLCTs was under the control of the P_tet_ promoter and induction was performed with 20 ng/mL anhydrotetracycline. The arrows indicate the two major bands corresponding to the HMW and LMW units of the SLCT-4 naturally present in the R20291 strain. The size of the bands varies depending on the SLCT. Note that the SLCT-11 LMW subunit is not migrating to the expected size because it is glycosylated. (M = MW marker).

**FIGURE 2.**
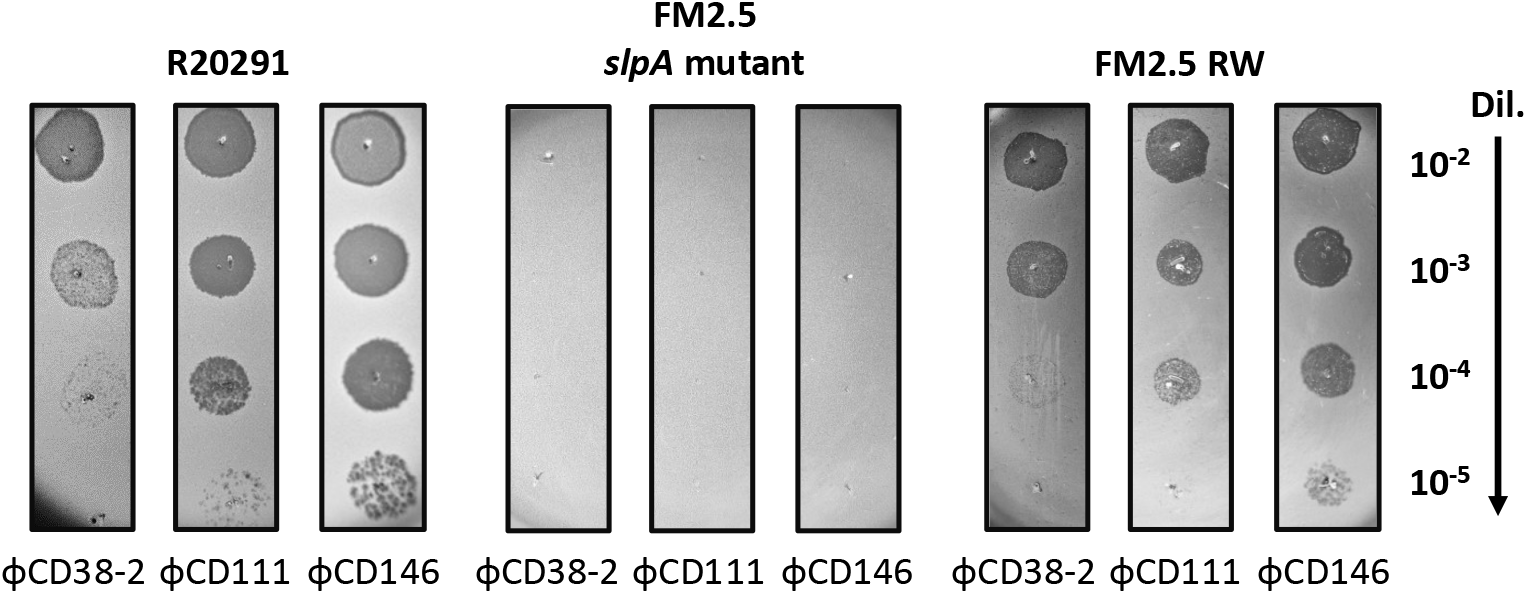
Spot test assays with the WT R20291, FM2.5 *slpA* mutant, and FM2.5RW watermarked revertant. Serial 10-fold dilutions of phages phiCD38-2, phiCD111 and phiCD146 were spotted on top of bacterial lawns of the indicated strains. Dark zones indicate bacterial lysis.

### The lack of SlpA impairs phage adsorption to host cells

The loss of phage susceptibility in absence of SlpA suggested that phage adsorption was impaired. To verify this, we performed phage adsorption assays and compared the FM2.5 mutant with the WT R20291 strain as a control. As shown in Fig. 3, the phages adsorbed to high levels onto the WT strain, with values of 92.9% ± 1.7% for phiCD38-2, 81.0% ± 6.7% for phiCD146 and 68.7% ± 10.8% for phiCD111. However, adsorption onto the FM2.5 mutant was severely affected, dropping to complete absence of adsorption for phiCD38-2 and phiCD111 and 22.5 ± 13.1% for phiCD146. In our experience, adsorption values below 50% did not result in productive infection. Together, these results further supported that SlpA is required by all three phages for infection, by allowing adsorption and close contact with the cell surface. Our results also suggest that in absence of SlpA, there is no alternative receptor involved.

**FIGURE 3.**
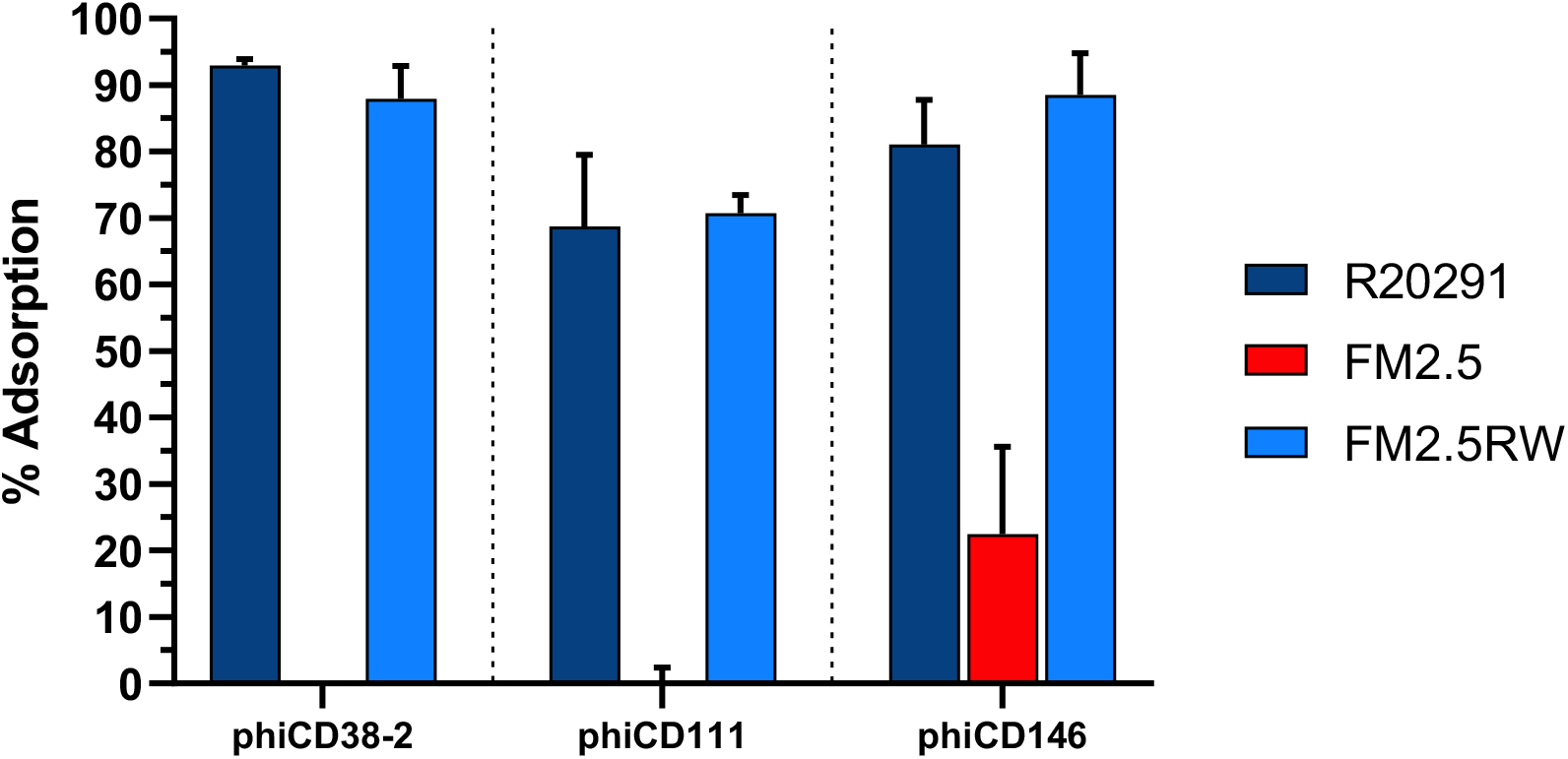
Phage adsorption assays on the WT R20291, FM2.5 *slpA* mutant, FM2.5RW watermarked revertant, and the FM2.5 *slpA* mutant complemented with various SLCTs. Data presented are the mean ± SEM of at least 3 replicates from a minimum of two independent experiments. Adsorption values for phiCD38-2 and phiCD111 on the FM2.5 mutant were below the limit of detection.

The three *Siphoviridae* phages tested are genetically related and share significant DNA homology. An all-against-all BLASTn analysis using Gegenees (40) showed that phiCD38-2 and phiCD146 share 79% whole genome identity, whereas phiCD38-2 and phiCD111 share 71% identity (data not shown). Whole proteome comparison using Phamerator (41) confirmed the extensive similarity between the proteome of the three phages, but also revealed variations in several proteins, including the tail fiber proteins gp20/21 and the predicted receptor binding protein (RBP) gp21/22 (Supplementary Fig. 1). These variations in tail fibers and RBP could possibly explain the different host ranges observed with these phages (7) and the different adsorption patterns observed on the WT strain, with phiCD111 adsorbing less than the other two phages (Fig. 3).

### *Siphoviridae* phages can use more than one SLCT as their receptor

Although the above results strongly support the role of SlpA as a phage receptor, we could not exclude a possible destabilization of the cell surface architecture caused by the lack of SlpA, which in turn could affect phage adsorption and prevent infection. To rule out this possibility, we complemented the FM2.5 mutant with one of 12 other SLCTs described previously (Table 1) (34, 42). Of note, Avidocin-CD have been shown to use more than one SLCT as their receptor to kill *C. difficile* (34). Hence, we sought to determine if phiCD38-2, phiCD111 and phiCD146 could also use alternative SLCTs to infect the R20291 strain, in addition to the natural SLCT-4.

**Table 1.** List of strains, plasmids, and phages used in this study.

| **Table 1.** List of strains, plasmids, and phages used in this study. |  |  |
| --- | --- | --- |
| **Strain, plasmids, or phages** | **Characteristic / description** | **Reference / source** |
| *Clostridium difficile* |  |  |
| R20291 | Epidemic isolate, ribotype 027 | (52) |
| R20291 FM2.5 | R20291 FM2.5 mutant carrying a G283A mutation in the *slpA* gene (SLCT-4) creating a premature stop codon (TAA) at position 289 | (34) |
| R20291 FM2.5RW | R20291 FM2.5 revertant carrying a functional watermarked copy of the *slpA* gene (SLCT-4) with two synonymous changes (A282G and A285T) | (34) |
| LCUS 1039 | R20291 FM2.5 *slpA*- mutant containing pJAK017 expressing SLCT-1 | This study |
| LCUS 1043 | R20291 FM2.5 *slpA*- containing pJAK023 expressing SLCT-2 under the control of the P<sub>tet</sub> inducible promoter | This study |
| LCUS 1040 | R20291 FM2.5 *slpA*- containing pJAK018 expressing SLCT-6 under the control of the P<sub>tet</sub> inducible promoter | This study |
| LCUS 1046 | R20291 FM2.5 *slpA*- containing pSEW027 expressing SLCT-7 under the control of the P<sub>tet</sub> inducible promoter | This study |
| LCUS 1041 | R20291 FM2.5 *slpA*- containing pJAK001 expressing SLCT-7b under the control of the P<sub>tet</sub> inducible promoter | This study |
| LCUS 1042 | R20291 FM2.5 *slpA*- containing pJAK019 expressing SLCT-8 under the control of the P<sub>tet</sub> inducible promoter | This study |
| LCUS 1371 | R20291 FM2.5 *slpA*- containing pJAK019.2 expressing SLCT-8 under the control of the P<sub>cwp2</sub> promoter | This study |
| LCUS 1047 | R20291 FM2.5 *slpA*- containing pJAK020 expressing SLCT-9 under the control of the P<sub>tet</sub> inducible promoter | This study |
| LCUS 1375 | R20291 FM2.5 *slpA*- containing pJAK020.2 expressing SLCT-9 under the control of the P<sub>cwp2</sub> promoter | This study |
| LCUS 1038 | R20291 FM2.5 *slpA*- containing pJAK003 expressing SLCT-10 under the control of the P<sub>tet</sub> inducible promoter | This study |
| LCUS 1048 | R20291 FM2.5 *slpA*- containing pAAM0013 expressing SLCT-11 under the control of the P<sub>tet</sub> inducible promoter | This study |
| LCUS 1374 | R20291 FM2.5 *slpA*- containing pAAM0013.2 expressing SLCT-11 under the control of the P<sub>cwp2</sub> promoter | This study |
| LCUS 1045 | R20291 FM2.5 *slpA*- containing pJAK021 expressing SLCT-12 under the control of the P<sub>tet</sub> inducible promoter | This study |
| LCUS 1044 | R20291 FM2.5 *slpA*- containing pJAK022 expressing SLCT-13 under the control of the P<sub>tet</sub> inducible promoter | This study |
| LCUS 1372 | R20291 FM2.5 *slpA*- containing pJAK022.2 expressing SLCT-13 under the control of the P<sub>cwp2</sub> promoter | This study |
| LCUS 1037 | R20291 FM2.5 *slpA*- containing pJAK002 expressing SLCT-H2/6 under the control of the P<sub>tet</sub> inducible promoter | This study |
| *Escherichia coli* |  |  |
| TOP10 | Cloning strain for pRPF plasmids | Invitrogen |
| CA434 | HB101 carrying plasmid R702 used for conjugation into *C. difficile* | (9) |
| Bacteriophages |  |  |
| phiCD38-2 | *Siphoviridae* | (44) |
| phiCD111 | *Siphoviridae* | (7) |
| phiCD146 | *Siphoviridae* | (7) |
| phiCD506 | *Myoviridae* | (7) |
| phiCD508 | *Myoviridae* | (7) |
| phiMMP02 | *Myoviridae* | (43) |
| phiMMP03 | *Myoviridae* | (7) |
| phiMMP04 | *Myoviridae* | (7) |
| **Plasmids** |  |  |
| pRPF144 | Reporter plasmid carrying the *gusA* gene under the control of the P<sub>cwp2</sub> constitutive promoter, and cloned into the BamHI and SacI sites. | (51) |
| pRPF185 | Reporter plasmid carrying the *gusA* gene under the control of the P<sub>tet</sub> inducible promoter, and cloned into the BamHI and SacI sites. | (51) |
| pJAK001 | pRPF185:SLCT-7b | (34) |
| pJAK002 | pRPF185:SLCT-H2/6 | (34) |
| pJAK003 | pRPF185:SLCT-10 | (34) |
| pJAK017 | pRPF185:SLCT-1 | (34) |
| pJAK018 | pRPF185:SLCT-6 | (34) |
| pJAK019 | pRPF185:SLCT-8 | (34) |
| pJAK019.2 | pRPF144:SLCT-8 | (34) |
| pJAK020 | pRPF185:SLCT-9 | (34) |
| pJAK020.2 | pRPF144:SLCT-9 | (34) |
| pJAK021 | pRPF185:SLCT-12 | (34) |
| pJAK022 | pRPF185:SLCT-13 | (34) |
| pJAK022.2 | pRPF144:SLCT-13 | (34) |
| pJAK023 | pRPF185:SLCT-2 | (34) |
| pSEW027 | pRPF185:SLCT-7 | (34) |
| pAAM0013 | pRPF185:SLCT-11 | (34) |
| pAAM0013.2 | pRPF144:SLCT-11 | (34) |

Representative members from each 12 SLCTs, cloned into the pRPF185 plasmid backbone, under the control of a P_tet_ tetracycline-inducible promoter, or in pRPF144 plasmid, under the control of the constitutive promoter P_cwp2_, were transferred by conjugation and expressed in the FM2.5 mutant. Correct expression of each SLCT was confirmed upon anhydrotetracycline induction by SDS-PAGE after glycine extraction (Fig. 1). The level of expression of the SLCTs cloned in front of the constitutive P_cwp2_ promoter were comparable to the induced forms (Supplementary Fig. 2)

Next, phage susceptibility assays were performed with all complemented strains and the three siphophages. Expression of the SLCT-6 restored susceptibility to phiCD111, and SLCT-11 rendered bacteria susceptible to infection by phiCD38-2 and phiCD146 (Fig. 4). All the other SLCT-complemented strains were found to be resistant to infection by the siphophages and no lysis zones were detected (data not shown, Table 2). The predicted RBPs (gp21) from phiCD38-2 and phiCD146 are 100% identical while they share 82% identity with gp22 from phiCD111 (Supplementary Fig. 1B). This might explain why phiCD111 did not infect cells expressing SLCT-11 and why only phiCD111 infected cells expressing SLCT-6. We cannot exclude that other tail proteins could be involved in receptor recognition as well. For instance, in the present case, the tail fiber protein gp20 from phiCD38-2 is different from gp20 from phiCD146 and from gp21 of phiCD111. These differences could possibly explain our observations, although further investigation will be required (Supplementary Fig. 1C). Nevertheless, our results show that the three *Siphoviridae* phages tested can use at least two SLCTs each to infect *C. difficile* and that this interaction is specific since not all SLCTs could restore susceptibility to infection.

**FIGURE 4.**
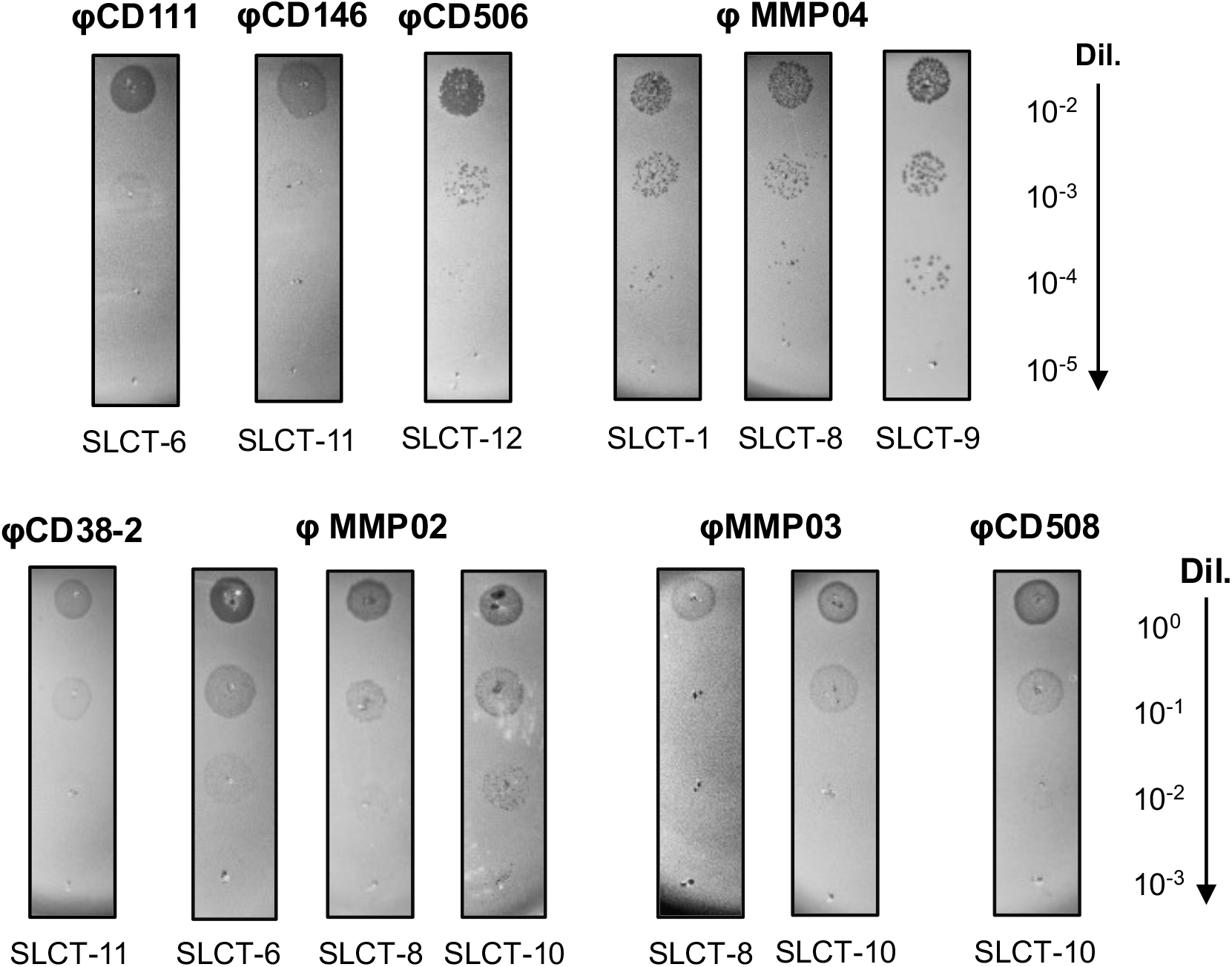
Phage susceptibility assays of the FM2.5 *slpA* mutant complemented with different SLCTs. Serial 10-fold dilutions of the indicated phages (titers of undiluted phage stocks = 10^9^ PFU/mL) were spotted on top of bacterial lawns of the indicated strains. Darker zones/plaques indicate bacterial lysis although extensive lysogeny was observed with some phages, making spots more difficult to observe. The experiments were repeated several times and only representative positive infections are shown here. Also, non-specific lysis was not observed in the absence of a productive infection since no lysis zone could be seen with other SLCTs using undiluted phage lysates.

**Table 2.**
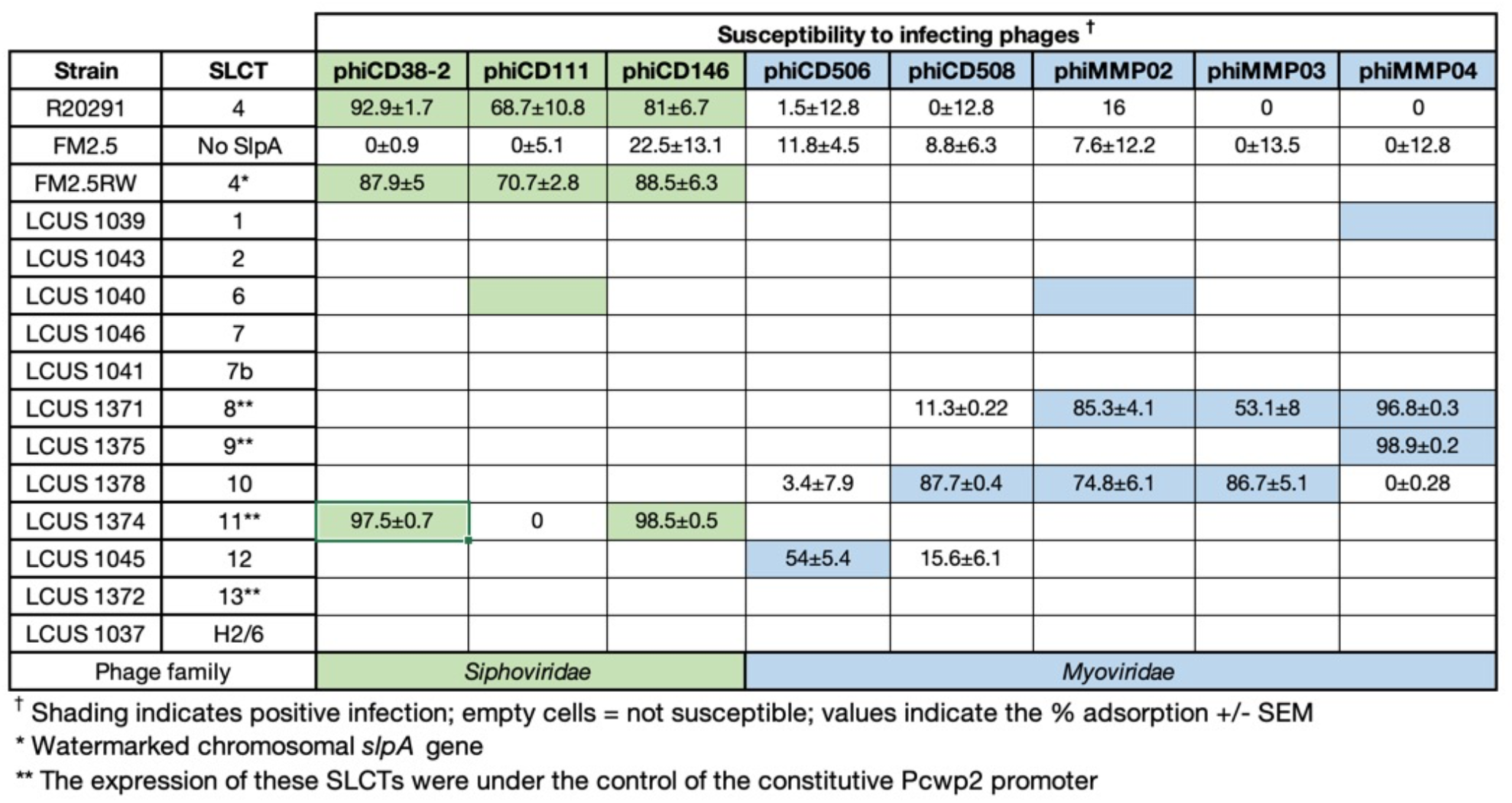
Susceptibility to phage infection of the FM2.5 mutant complemented with various SLCTs.

### Expression of specific SLCTs into the FM2.5 mutant confers susceptibility to additional phages

Besides the *Siphoviridae* phages phiCD38-2, phiCD111 and phiCD146, no other phages from our collection can form plaques on the WT R20291. The most likely reason to explain the lack of infection by some phages is the absence of a suitable host receptor on the bacterial surface.

Different *Myoviridae* phages from our collection were used in spot test assays against the FM2.5 mutant carrying one of the 12 SLCTs expressed from a plasmid. As shown in Table 2 and Fig. 4, the complemented mutant became susceptible to infection by phiCD506, phiCD508, phiMMP02, phiMMP03 and phiMMP04 when one of SLCT-1, −6, −8, −9, 10, or −12 was expressed. These results confirm that SlpA is also used as a receptor by several *Myoviridae* phages, thus corroborating previous studies (34, 38, 39). Of note, three of the *Myoviridae* phages also recognized more than one SLCT. Also, except for phiMMP02 and phiCD111 that both recognized SLCT-6, *Myoviridae* and *Siphoviridae* phages seemed to bind different SLCTs, and no overlap was observed in susceptibility testing. We performed phage adsorption assays to investigate the attachment of the phages depending on the SLCT expressed. As shown Table 2, high adsorption ratios were observed for phages that were able to infect a given complemented strain, but little to no adsorption was observed when no infection occurred. Although we did not test adsorption with all possible combinations of phages and SLCTs, a minimum of ~50% adsorption was found to be necessary to allow infection in our assays, while little to no adsorption correlated with the absence of infection.

The phages used in this study have been originally isolated on four different strains (7, 43, 44). Strain LCUS0274 is a ribotype 027 strain on which phiCD38-2, phiCD111 and phiCD146 were isolated and propagated, that expresses SLCT-4 like the R20291 strain in this study. Likewise, phages phiCD508, phiMMP02 and phiMMP03 were isolated on strain LCUS0117 (7, 43) that expresses SLCT-10. Accordingly, these phages recognized SLCT-10 in the FM2.5 complemented strain (Table 2). In summary, different *Myoviridae* phages can use one or more SLCTs as their receptor. These results clearly demonstrate that SlpA is a general receptor used by many *C. difficile* phages.

### Relationship between SLCT, phage RBP and susceptibility to infection

We sought to determine if some relationship could be made between the SLCT, the phage RBP and the observed susceptibility to infection. Multiple alignments were built using the SlpA protein on one hand and the predicted phage RBPs on the other (Supplementary Fig. 3 and 4). Phylogenetic trees were generated and then compared with the susceptibility profile of the different complemented strains (Fig. 5). Although some of the RBPs are highly similar (e.g. phiCD146 vs. phiCD111 or phiCD508 vs. phiMMP03), slight differences in host range were observed. This suggests that some of the diverging regions between these RBPs are involved in binding with these specific SlpA isoforms or that other phage proteins are involved in binding, such as the tail fiber gp20/21 (Supplementary Fig. 1). Regarding the *Myoviridae* phages, phiMMP02 and phiMMP04 both recognized SLCT-8, and although they were grouped together in the phylogenetic tree based on their RBP, the level of amino acid conservation is only 67% over 38% of the protein. On the other hand, phiCD508 and phiMMP03 recognized SLCT-10 and their RBP are closely related and share 92% amino acid identity over 100% of the protein. Yet, SLCT-8 is not bound by phiCD508 whereas it is by phiMMP03. In addition, phiMMP02, and phiMMP03 both recognized SLCT-8 and SLCT-10, but they share only 23% sequence identity over 50% of the protein. These observations make prediction of the phage host range based on the RBP alone very difficult. Note that we also did a clustering analysis using only the LMW fragment of the SLCT to generate the phylogenetic tree and very similar results were obtained (Supplementary Fig. 4 and 5). In summary, although we could observe tendencies, larger analyses will be necessary to draw conclusions linking a particular RBP with specific SLCTs.

**FIGURE 5.**
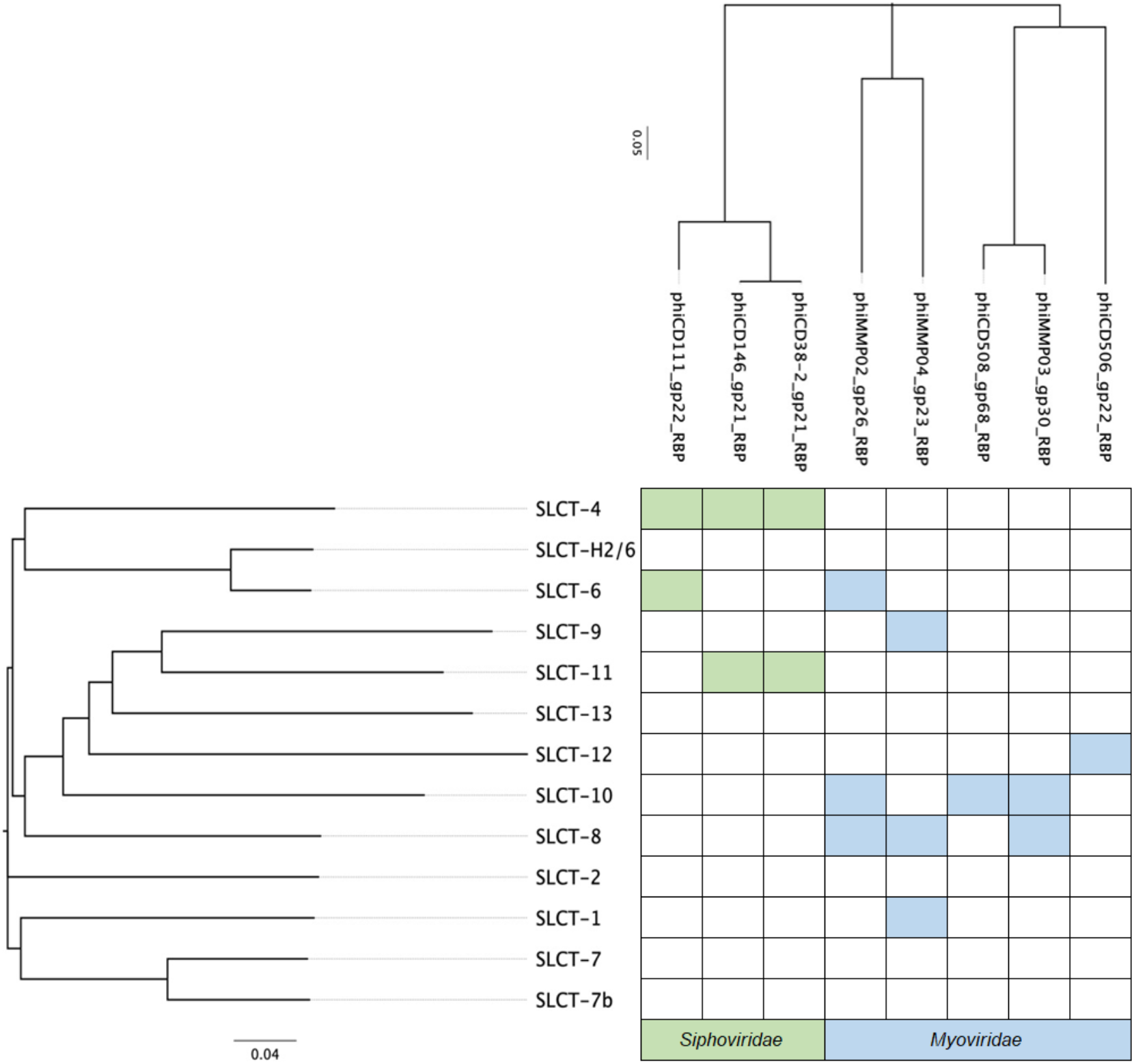
Susceptibility to phage infection in function of SLCT and RBP phylogeny. The full-length SlpA amino acid sequence was aligned, and a phylogenetic tree was constructed, creating 4 different clades (indicated by colored lines). The amino acid sequences of the predicted phage RBPs were also aligned, and a phylogenetic tree was built. The branches of both trees were reorganized in function of the susceptibility to phage infection (shaded cells).

## DISCUSSION

The identification of the receptor(s) used by phages to infect *C. difficile* has been a topic of great interest for many years. Here we used a panel of 3 *Siphoviridae* and 5 *Myoviridae* phages in a single bacterial model based on the FM2.5 *slpA* mutant isolated previously (34) to provide unambiguous experimental proof that many phages of *C. difficile* use the cell surface protein SlpA as their receptor to initiate infection. Previous work on Avidocin-CD provided indirect evidence that *C. difficile* myophages could use SlpA as their host receptor (34). More recent reports brought additional evidence that the S-layer acts as the receptor for two *C. difficile* myophages (φCD1801 and φHN10) (36, 39) and a siphophage (phiCDHS-1) (38). However, the study with φHN10 involved phage binding assays in gel retardation assays with crude cell wall extracts, and no direct evidence could be provided showing that SlpA was the receptor (36). The report by Whittle *et al* involved adsorption assays with the myophage φCD1801 and the *C. difficile* strain 630 co-expressing one of three different SLCTs, in addition to its native S-layer. However, the authors did not confirm expression of the exogenous SlpA gene and therefore could not completely exclude the possibility that an indirect effect could be responsible for the observed adsorption results. Also, the absence of infection in the strain 630 background did not allow to conclude that SlpA can be used by phages as a receptor to infect cells and that adsorption was not due to other factors (39). Finally, the recent report on the siphophage phiCDHS-1 showed that the absence of an S-layer in the R20291 FM2.5 mutant prevented phage infection, suggesting that SlpA is the receptor used by this phage. However, in the absence of complementation and adsorption assays, it could not be fully concluded that SlpA is really the receptor used by phiCDHS-1 (38).

Because all our experiments were conducted in the same genetic background and SlpA isoforms were expressed only one at a time in complementation assays, we could readily compare the infection efficacy of our phages in function of the SLCT expressed. While this approach alleviated several of the limitations that occur during phage host range assays using multiple bacterial strains, the level of expression of SlpA from a plasmid was generally lower than the normal chromosomal allele. Except for SLCT-8 that was expressed to levels close to the natural SLCT-4, all the other SLCTs were expressed to lower levels, even when using the constitutive promoter P_cwp2_. Consequently, many of the positive infections observed with some SLCTs were weaker than what is normally observed with the natural host of the corresponding phages. Nevertheless, we can exclude a problem in the expression of SlpA since adsorption was high in most cases of positive infection and adsorption values were very similar to those observed on the natural replicating host of the corresponding phages. For example, phiCD38-2 and phiCD146 adsorbed to very high levels on the SLCT-11, but infected cells less efficiently than the WT R20291. Also, we could see strong infection with phiMMP04 on SLCT-1, −8 and −9, even if SLCT-1 was not expressed as efficiently compared with the other two SLCTs. A previous study by Thanki *et al* (45) suggested that high adsorption (>80%) was required for productive infection, however, we could still detect productive infection with certain phages even if the adsorption was only slightly above 50% (e.g. phiMMP03 on SLCT-8 and phiCD506 on SLCT-12). Therefore, the differences observed in the efficacy of infection might be related to other factors than adsorption per se. Supporting this, our three siphophages can infect the WT R20291 strain very well, so we can rule out the possibility that the reduced efficacy of infection observed with phiCD38-2 on SLCT-11 was due to problems in replication of the phages in this strain. One possible explanation could be that the interaction between the RBP and different SLCTs is not always optimal despite strong adsorption. The process of baseplate docking to the receptor might also be compromised. Conformational changes in the baseplate structure, as seen in the lactococcal phage p2 (18, 19), might also be required for infection but compromised with certain SLCTs.

It is noteworthy to mention that 8 SLCTs out of the 13 SLCTs tested could be targeted by phages in our study. The absence of infection with 5 SLCTs could simply reflect the small phage collection used herein. Nevertheless, an interesting observation we made was that the SLCTs recognized by *Siphoviridae* and *Myoviridae* were different, except for SLCT-6, suggesting that the two phage families have different binding patterns. Also, SLCT-8 and SLCT-10 were recognized by multiple *Myoviridae* phages with diverse RBPs. These SlpA isoforms are not closely related, so it will be interesting to investigate if specific domains, structures, or amino acid residues could explain this greater binding capacity. Our clustering analysis did not allow us to find any clear link between SLCTs, RBPs and susceptibility to infection. Similar conclusions were reported whereby phages with identical tail fibers displayed different host ranges (45). These elements argue in favor of a complex interaction that would involve the RBP in addition to other tail components. Future studies with a more diverse panel of well-characterized phages will be required to get a refined view of the interaction between phages and the S-layer.

Phage therapy has gained interest in recent years and treatment of *C. difficile* infections with therapeutic phages could have huge advantages over currently available antibiotics. Therefore, phage therapy has been viewed as a potential alternative to fight *C. difficile* infections (3, 4). However, it is currently constrained by the fact that all phages known to infect *C. difficile* are temperate, and thus, capable of lysogeny (6). Using these phages for therapy is not recommendable, as exemplified by several reports showing growth rebound and lysogeny after treatment, particularly when single phages were used (6, 46–49). Fortunately, the issue of lysogeny can be circumvented by genetic engineering (49), but another important limitation is the relatively narrow host range of available phages. Even with phage cocktails, it is difficult to target multiple strains (3). Here, experimental demonstration that SlpA is a general host receptor used by many *C. difficile* phages opens the way to genetic engineering of tail genes to target multiple strains with a limited number of phages. One could argue that mutation of the SlpA protein would quickly lead to phage resistance, which is true, and the FM2.5 and FM2.6 mutants confirm this possibility. However, it was shown that the lack of an S-layer comes with a huge fitness and virulence cost (34). Therefore, development of phage resistance through SlpA mutation would be deleterious for phage therapy applications, but would also strongly impair *C. difficile* capacity to cause disease.

In summary, our study brings unambiguous experimental proof that SlpA is a receptor used by both *Myoviridae* and *Siphoviridae* phages and therefore seems to be a general phage receptor. A lot remains to be done to fully understand how *C. difficile* phages interact with the S-layer, how the RBP binds to SlpA and whether accessory proteins are involved or if the adsorption and/or DNA injection mechanisms varies between phage subgroups. Ongoing structural work that aims at reconstructing phage architecture will clearly bring insightful knowledge in this regard.

## MATERIALS AND METHODS

### Bacterial strains, plasmids, and bacteriophages

A comprehensive list of bacterial strains, plasmids, and phages used in this work is presented in Table 1. *C. difficile* was grown in an anaerobic chamber (Coy Laboratories, anaerobic conditions H_2_ 10%, CO_2_ 5%, and N_2_ 85%) at 37°C in pre-reduced brain heart infusion (BHI) broth or TY broth (2% yeast extract, 3% tryptose, pH 7.4). Thiamphenicol (15 μg mL) or norfloxacin (12 μg/mL) were added when necessary. *Escherichia coli* was grown in aerobic conditions in Luria–Bertani (LB) broth in an incubator with agitation at 37°C with, chloramphenicol (25 μg/mL) or kanamycin (50 μg/mL), when necessary.

### Bacteriophage amplification and titration

Phage lysates (>10^9^ PFU/mL) were prepared in TY broth using standard phage amplification protocols and titers were determined using the soft agar overlay method, as described previously (50). Phage lysates were filtered through 0.45 μm membranes and stored at 4°C. Phage titers were verified regularly.

### Conjugation of plasmid DNA into *C. difficile*

Plasmids carrying the different SLCTs described previously were transferred by conjugation into the FM2.5 mutant strain as previously described using the *Escherichia coli* CA434 strain as the donor (Table 1) (10).

### Induction of *slpA* expression in *C. difficile*

The plasmids containing one of the 12 SlpA isoforms cloned into the pRPF185 plasmid were under the control of the inducible P_tet_ promoter (34, 51). To induce the expression of SlpA, bacteria were grown in 10 mL of TY broth up to an OD_600nm_ = 0.4 and anhydrotetracycline was added to a final concentration of 20 ng/mL. Cultures were grown overnight and used to assess susceptibility to phage infection, and to assess the level of SlpA expression by SDS-PAGE following glycine extraction (see below).

### Subcloning of SLCTs for constitutive expression in *C. difficile*

The SLCT 8, 9, 11 and 13 were subcloned into the pRPF144 plasmid to allow constitutive expression of the *slpA* gene under control of the P_cwp2_ promoter. To do this, the pJAK019, pJAK020, pJAK022 and pAAM013 plasmids (Table 1) were digested with BamHI and SacI restriction enzymes to excise the *slpA* gene. The pRPF144 plasmid was also digested with BamHI and SacI to remove the *gusA* gene (51). Digested fragments were extracted from a 0,8% agarose gel using the QIAquick Gel Extraction Kit (QIAgen, Mississauga, ON). The *slpA* gene fragments were then ligated into the pRPF144 backbone using T4 DNA ligase at room temperature for 1 h and the ligation products were transformed in *E. coli* MC1061 using standard protocols. The resulting plasmids were then purified using the Geneaid High-Speed Mini Kit (Froggabio, Concord, ON) and then sequenced at the Université Laval sequencing center. The validated plasmids were then transformed into *E. coli* CA434 donor cells and conjugated into the *C. difficile* FM2.5 strain as described above.

### Glycine extraction of cell surface proteins

We assessed the expression of the SlpA proteins on SDS-PAGE after performing a surface protein extraction of the induced cultures. The samples were prepared as follows: 10 mL of the induced *C. difficile* cultures were centrifugated at 4,000xg followed by a washing step with PBS 1X. We suspended the pellet with 200 μL of 0.2M glycine, pH 2.2 and we incubated at room temperature for 30 minutes. We centrifugated again during 5 minutes at 10,000xg. Then we transferred 150 μL of the supernatant into a new tube, and we adjusted the pH to 7.5 using a solution of Tris HCl 2M, pH 7.5. We mixed 15 μL of the samples and 5 μl of loading buffer 4X (Tris-HCl 200mM pH 6.8, DTT 400mM, 8% SDS, 0.4% bromophenol blue and 40% glycerol). We then ran the samples 12% polyacrylamide gels (BioShop). The migration of the sample was performed in a Mini-Protean^®^ Tetra Cell apparatus (Bio-rad, Mississauga, ON, Canada), using a voltage of 100V for 25 min, followed by 1h at 150V. The gels were stained with Coomassie blue.

### Phage susceptibility testing on FM2.5 SLCT-complemented strains

Spot tests using a standard soft agar overlay method were used to determine susceptibility to phage infection of different FM2.5 complemented strains (50). The night before the experiment, a preculture of a complemented strain was inoculated in 5 mL of TY broth supplemented with thiamphenicol and incubated at 37°C under anaerobic conditions. The next day, a fresh 5-10 mL culture was inoculated with 5% of the O/N preculture, and thiamphenicol was added, as well as 20 ng/mL anhydrotetracycline. The OD at 600nm was then monitored regularly. Simultaneously, we prepared TY with 0.3% agarose maintained at a temperature of 55°C. When the bacterial culture reached an OD_600nm_ of 0.8, 4 mL of soft agarose was mixed with antibiotics when necessary (thiamphenicol 15 μg/mL and anhydrotetracycline 20 ng/mL), 0.67 mL of the bacterial culture and salts (100mM MgCl_2_ and 0,3mM CaCl_2_). We then poured the tube content over square Petri dishes containing TY bottom agar (1% agar) with antibiotics when required (thiamphenicol 15 μg/mL and anhydrotetracycline 20ng/mL). Once the top agarose has hardened, 5 μL of serially diluted phage lysates were deposited directly on top of the soft agarose overlay. We incubated overnight at 37°C in an anaerobic chamber. Zones of lysis in the bacterial lawn revealed a productive phage infection.

### Bacteriophage adsorption assays

Phage adsorption assays were performed as described previously (10). Briefly, bacteria from an overnight culture were grown in TY broth for 24 h. Then, 0.9 mL of culture was mixed with 1×10^5^ PFU of the desired phage, in the presence of salts (10 mM each of MgCl_2_ and CaCl_2_), in a final volume of 1 mL. Phages were allowed to adsorb for 30 minutes at 37°C. Cells were then collected by centrifugation at 13,000g for 1 minutes. Free phages in the supernatant that did not adsorb were enumerated after serial 10-fold dilutions on standard soft agarose overlays as described earlier and titers were analyzed against the initial phage input. The adsorption ratio was calculated using the following formula: 100 – ([residual titer/ initial titer] × 100). Each experiment was done in technical replicate. Plots were generated using the mean adsorption ± standard error of the mean (SEM).

## ACKNOWLEDGMENTS

All authors contributed to the manuscript. ALMR, AAU, MMA, JRG, and MOB performed experiments, analyzed data and contributed to draft writing. JK, RF and GG contributed material and reviewed the draft. OS reviewed the draft. LCF designed the experiments, analyzed data, drafted portions of the manuscript, and completed the final version. All authors agreed with the final version. This research was funded by an NSERC discovery grant (RGPIN RGPIN-2020-05776 and RGPIN-2015-06334). MOB received a Gérard-Eugène Plante master scholarship. MMA and AAU received a master scholarship from the faculty of medicine and health sciences of the UdeS, JRG received an NSERC PhD scholarship. LCF is a member of the Centre de recherche du Centre hospitalier universitaire de Sherbrooke (CRCHUS).

## REFERENCES

1. Lin DM, Koskella B, Lin HC. 2017. Phage therapy: An alternative to antibiotics in the age of multidrug resistance. World J Gastrointest Pharmacol Ther 8:162–173.

2. Guery B, Galperine T, Barbut F. 2019. *Clostridioides difficile*: diagnosis and treatments. Bmj 366:l4609.

3. Nale JY, Spencer J, Hargreaves KR, Buckley AM, Trzepiński P, Douce GR, Clokie MRJ. 2016. Bacteriophage Combinations Significantly Reduce *Clostridium difficile* Growth In Vitro and Proliferation In Vivo. Antimicrobial agents and chemotherapy 60:968–981.

4. Venhorst J, Vossen JMBM van der, Agamennone V. 2022. Battling Enteropathogenic Clostridia: Phage Therapy for *Clostridioides difficile* and *Clostridium perfringens*. Front Microbiol 13:891790.

5. Abt MC, McKenney PT, Pamer EG. 2016. *Clostridium difficile* colitis: pathogenesis and host defence. Nat Rev Microbiol 14:609–620.

6. Heuler J, Fortier L-C, Sun X. 2021. *Clostridioides difficile* phage biology and application. Fems Microbiol Rev 45:fuab012.

7. Sekulovic O, Garneau JR, Néron A, Fortier L-C. 2014. Characterization of Temperate Phages Infecting *Clostridium difficile* Isolates of Human and Animal Origins. Appl Environ Microb 80:2555–2563.

8. Boudry P, Semenova E, Monot M, Datsenko KA, Lopatina A, Sekulovic O, Ospina-Bedoya M, Fortier L-C, Severinov K, Dupuy B, Soutourina O. 2015. Function of the CRISPR-Cas System of the Human Pathogen *Clostridium difficile*. Mbio 6:e01112–15.

9. Purdy D, O’Keeffe TAT, Elmore M, Herbert M, McLeod A, Bokori-Brown M, Ostrowski A, Minton NP. 2002. Conjugative transfer of clostridial shuttle vectors from Escherichia coli to *Clostridium difficile* through circumvention of the restriction barrier. Mol Microbiol 46:439–452.

10. Sekulovic O, Bedoya MO, Fivian-Hughes AS, Fairweather NF, Fortier L. 2015. The *Clostridium difficile* cell wall protein CwpV confers phase-variable phage resistance. Mol Microbiol 98:329–342.

11. Fortier L-C. 2017. Chapter Five The Contribution of Bacteriophages to the Biology and Virulence of Pathogenic Clostridia. Adv Appl Microbiol 101:169–200.

12. Labrie SJ, Samson JE, Moineau S. 2010. Bacteriophage resistance mechanisms. Nat Rev Microbiol 8:317–327.

13. Dowah ASA, Clokie MRJ. 2018. Review of the nature, diversity and structure of bacteriophage receptor binding proteins that target Gram-positive bacteria. Biophysical Rev 10:535–542.

14. Taylor NMI, Prokhorov NS, Guerrero-Ferreira RC, Shneider MM, Browning C, Goldie KN, Stahlberg H, Leiman PG. 2016. Structure of the T4 baseplate and its function in triggering sheath contraction. Nature 533:346–352.

15. Ferreira RCG, Hupfeld M, Nazarov S, Taylor NM, Shneider MM, Obbineni JM, Loessner MJ, Ishikawa T, Klumpp J, Leiman PG. 2018. Structure and transformation of bacteriophage A511 baseplate and tail upon infection of *Listeria* cells. The EMBO journal 38:882–20.

16. Casjens SR, Hendrix RW. 2015. Bacteriophage lambda: Early pioneer and still relevant. Virology 479:310–330.

17. Vinga I, Baptista C, Auzat I, Petipas I, Lurz R, Tavares P, Santos MA, São-José C. 2012. Role of bacteriophage SPP1 tail spike protein gp21 on host cell receptor binding and trigger of phage DNA ejection. Molecular Microbiology 83:289–303.

18. Bebeacua C, Tremblay D, Farenc C, Chapot-Chartier M-P, Sadovskaya I, Heel M van, Veesler D, Moineau S, Cambillau C. 2013. Structure, adsorption to host, and infection mechanism of virulent lactococcal phage p2. Journal of virology 87:12302–12312.

19. Sciara G, Bebeacua C, Bron P, Tremblay D, Ortiz-Lombardia M, Lichière J, Heel M van, Campanacci V, Moineau S, Cambillau C. 2010. Structure of lactococcal phage p2 baseplate and its mechanism of activation. Proc National Acad Sci 107:6852–6857.

20. Dunne M, Hupfeld M, Klumpp J, Loessner MJ. 2018. Molecular Basis of Bacterial Host Interactions by Gram-Positive Targeting Bacteriophages. Viruses 10:397.

21. Monteville MR, Ardestani B, Geller BL. 1994. Lactococcal Bacteriophages Require a Host Cell Wall Carbohydrate and a Plasma Membrane Protein for Adsorption and Ejection of DNA. Appl Environ Microb 60:3204–3211.

22. Azam AH, Hoshiga F, Takeuchi I, Miyanaga K, Tanji Y. 2018. Analysis of phage resistance in *Staphylococcus aureus* SA003 reveals different binding mechanisms for the closely related Twort-like phages φSA012 and φSA039. Appl Microbiol Biot 102:8963–8977.

23. Letellier L, Boulanger P, Plançon L, Jacquot P, Santamaria M. 2004. Main features on tailed phage, host recognition and DNA uptake. Front Biosci 9:1228.

24. Silva JB, Storms Z, Sauvageau D. 2016. Host receptors for bacteriophage adsorption. Fems Microbiol Lett 363:fnw002.

25. Chaturongakul S, Ounjai P. 2014. Phage–host interplay: examples from tailed phages and Gram-negative bacterial pathogens. Front Microbiol 5:442.

26. Lanzoni-Mangutchi P, Banerji O, Wilson J, Barwinska-Sendra A, Kirk JA, Vaz F, O’Beirne S, Baslé A, Omari KE, Wagner A, Fairweather NF, Douce GR, Bullough PA, Fagan RP, Salgado PS. 2022. Structure and assembly of the S-layer in *C. difficile*. Nat Commun 13:970.

27. Fagan RP, Fairweather NF. 2014. Biogenesis and functions of bacterial S-layers. Nat Rev Microbiol 12:211–222.

28. Merrigan MM, Venugopal A, Roxas JL, Anwar F, Mallozzi MJ, Roxas BAP, Gerding DN, Viswanathan VK, Vedantam G. 2013. Surface-Layer Protein A (SlpA) Is a Major Contributor to Host-Cell Adherence of *Clostridium difficile*. Plos One 8:e78404.

29. O’Beirne S, Kirk JA, Fagan RP. 2019. *Clostridium difficile:* cell surface biogenesis. Access Microbiol 1.

30. Kirk JA, Banerji O, Fagan RP. 2017. Characteristics of the *Clostridium difficile* cell envelope and its importance in therapeutics. Microb Biotechnol 10:76–90.

31. Eidhin DN, Ryan AW, Doyle RM, Walsh JB, Kelleher D. 2006. Sequence and phylogenetic analysis of the gene for surface layer protein, *slpA*, from 14 PCR ribotypes of *Clostridium difficile*. J Med Microbiol 55:69–83.

32. Fagan RP, Albesa-Jové D, Qazi O, Svergun DI, Brown KA, Fairweather NF. 2009. Structural insights into the molecular organization of the S-layer from *Clostridium difficile*. Mol Microbiol 71:1308–1322.

33. Gebhart D, Williams SR, Bishop-Lilly KA, Govoni GR, Willner KM, Butani A, Sozhamannan S, Martin D, Fortier L-C, Scholl D. 2012. Novel High-Molecular-Weight, R-Type Bacteriocins of *Clostridium difficile*. J Bacteriol 194:6240–6247.

34. Kirk JA, Gebhart D, Buckley AM, Lok S, Scholl D, Douce GR, Govoni GR, Fagan RP. 2017. New class of precision antimicrobials redefines role of *Clostridium difficile* S-layer in virulence and viability. Sci Transl Med 9:eaah6813.

35. Gebhart D, Lok S, Clare S, Tomas M, Stares M, Scholl D, Donskey CJ, Lawley TD, Govoni GR. 2015. A Modified R-Type Bacteriocin Specifically Targeting *Clostridium difficile* Prevents Colonization of Mice without Affecting Gut Microbiota Diversity. Mbio 6:e02368–14.

36. Phothichaisri W, Ounjai P, Phetruen T, Janvilisri T, Khunrae P, Singhakaew S, Wangroongsarb P, Chankhamhaengdecha S. 2018. Characterization of Bacteriophages Infecting Clinical Isolates of *Clostridium difficile*. Front Microbiol 9:66–13.

37. Whittle MJ, Bilverstone TW, Esveld RJ van, Lücke AC, Lister MM, Kuehne SA, Minton NP. 2022. A Novel Bacteriophage with Broad Host Range against *Clostridioides difficile* Ribotype 078 Supports SlpA as the Likely Phage Receptor. Microbiol Spectr 10:e02295–21.

38. Dowah ASA, Xia G, Ali AAK, Thanki AM, Shan J, Millard A, Petersen B, Sicheritz-Pontén T, Wallis R, Clokie MRJ. 2021. The structurome of a *Clostridium difficile* phage and the remarkable accurate prediction of its novel phage receptor-binding protein. Biorxiv 2021.07.05.451159.

39. Whittle MJ, Bilverstone TW, Esveld RJ van, Lücke AC, Lister MM, Kuehne SA, Minton NP. 2022. A Novel Bacteriophage with Broad Host Range against *Clostridioides difficile* Ribotype 078 Supports SlpA as the Likely Phage Receptor. Microbiol Spectr 10:e02295–21.

40. Agren J, Sundström A, Håfström T, Segerman B. 2012. Gegenees: fragmented alignment of multiple genomes for determining phylogenomic distances and genetic signatures unique for specified target groups. PLoS One 7:e39107.

41. Gao R, Naushad S, Moineau S, Levesque R, Goodridge L, Ogunremi D. 2020. Comparative genomic analysis of 142 bacteriophages infecting Salmonella enterica subsp. enterica. Bmc Genomics 21:374.

42. Dingle KE, Didelot X, Ansari MA, Eyre DW, Vaughan A, Griffiths D, Ip CLC, Batty EM, Golubchik T, Bowden R, Jolley KA, Hood DW, Fawley WN, Walker AS, Peto TE, Wilcox MH, Crook DW. 2013. Recombinational Switching of the *Clostridium difficile* S-Layer and a Novel Glycosylation Gene Cluster Revealed by Large-Scale Whole-Genome Sequencing. J Infect Dis 207:675–686.

43. Meessen-Pinard M, Sekulovic O, Fortier L-C. 2012. Evidence of In Vivo Prophage Induction during *Clostridium difficile* Infection. Appl Environ Microb 78:7662–7670.

44. Sekulovic O, Meessen-Pinard M, Fortier L-C. 2011. Prophage-Stimulated Toxin Production in *Clostridium difficile* NAP1/027 Lysogens. J Bacteriol 193:2726–2734.

45. Thanki AM, Taylor-Joyce G, Dowah A, Nale JY, Malik D, Clokie MRJ. 2018. Unravelling the Links between Phage Adsorption and Successful Infection in *Clostridium difficile*. Viruses 10:411.

46. Meader E, Mayer MJ, Steverding D, Carding SR, Narbad A. 2013. Evaluation of bacteriophage therapy to control *Clostridium difficile* and toxin production in an in vitro human colon model system. Anaerobe 22:25–30.

47. Revathi G, Fralick JA, Rolfe RD. 2011. In vivo lysogenization of a *Clostridium difficile* bacteriophage ΦCD119. Anaerobe 17:125–129.

48. Meader E, Mayer MJ, Gasson MJ, Steverding D, Carding SR, Narbad A. 2010. Bacteriophage treatment significantly reduces viable *Clostridium difficile* and prevents toxin production in an in vitro model system. Anaerobe 16:549–554.

49. Selle K, Fletcher JR, Tuson H, Schmitt DS, McMillan L, Vridhambal GS, Rivera AJ, Montgomery SA, Fortier L-C, Barrangou R, Theriot CM, Ousterout DG. 2020. In Vivo Targeting of *Clostridioides difficile* Using Phage-Delivered CRISPR-Cas3 Antimicrobials. Mbio 11:e00019–20.

50. Sekulović O, Fortier L-C. 2016. *Clostridium difficile*, Methods and Protocols, p. 143–165. In Characterization of Functional Prophages in Clostridium difficile.

51. Fagan RP, Fairweather NF. 2011. *Clostridium difficile* Has Two Parallel and Essential Sec Secretion Systems*. J Biol Chem 286:27483–27493.

52. Stabler RA, He M, Dawson L, Martin M, Valiente E, Corton C, Lawley TD, Sebaihia M, Quail MA, Rose G, Gerding DN, Gibert M, Popoff MR, Parkhill J, Dougan G, Wren BW. 2009. Comparative genome and phenotypic analysis of *Clostridium difficile* 027 strains provides insight into the evolution of a hypervirulent bacterium. Genome Biology 10:R102.

