## Supplemental figures 1-5 for "The *Clostridioides difficile* S-Layer Protein A (SlpA) serves as a general phage receptor"

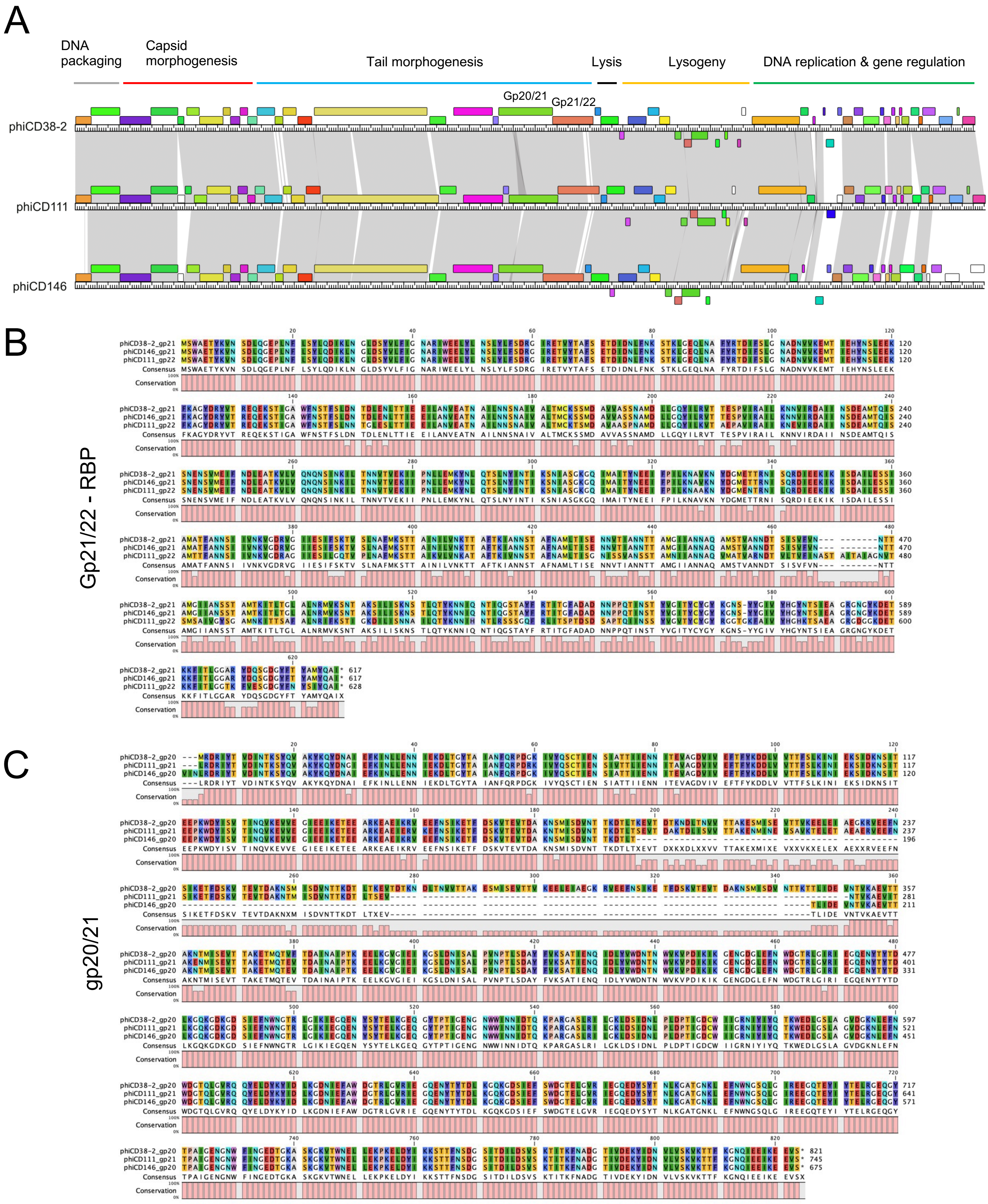


**Supplementary figure 1.** A) Genome-wide proteome comparison of the *Siphoviridae* phiCD38-2, phiCD111 and phiCD146 based on Phamerator analysis (24). Each linear map represents the proteome of a whole phage, and each colored box represents a protein of the same “phamily”. B) Alignment of the predicted RBP gp 21/22, and C) alignment of the predicted tail fiber gp20/21.

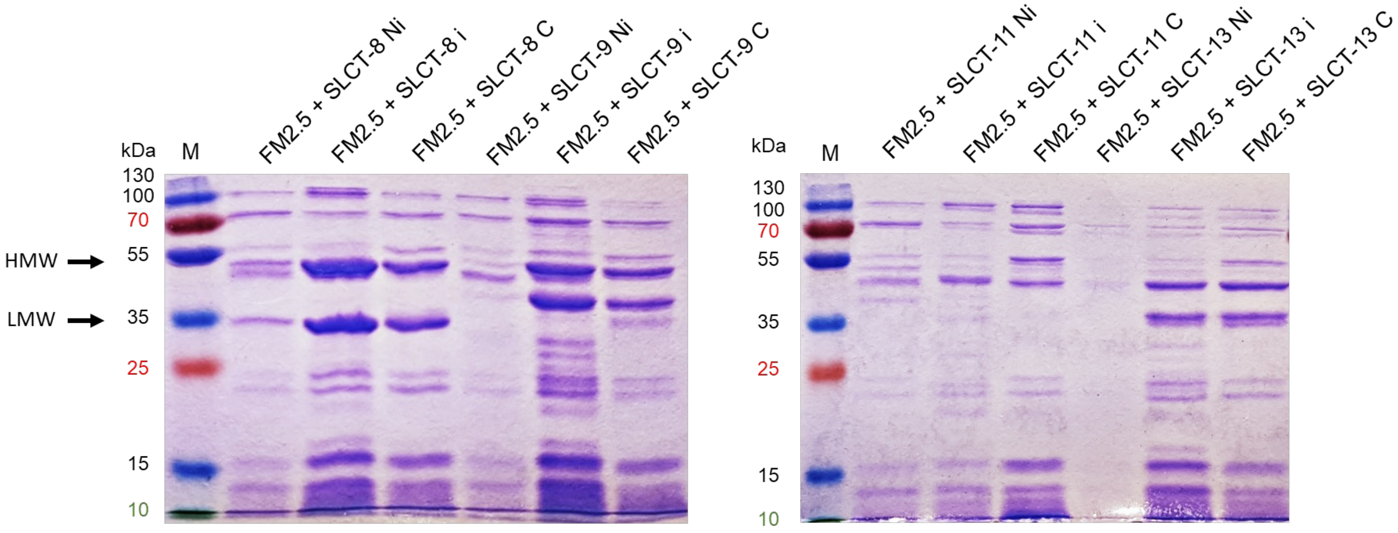


**Supplementary figure 2. Comparison between non-induced, induced, and constitutive expression of different SlpA isoforms.** Coomassie-stained 12% SDS-PAGE showing glycine extractions of surface proteins from FM2.5 mutant strains complemented with plasmids encoding SLCT-8, SLCT-9, SLCT-11 or SLCT-13. These isoforms were under the control of the $P_{tet}$ tetracycline inducible promoter or the $P_{cwp2}$ constitutive promoter. Ni = non-induced; i = induced with 20ng/mL anhydrotetracycline; C = constitutive. The arrows indicate the two major bands corresponding to the HMW and LMW fragments from SLCT-8. The size of the bands varies depending on the isoform. (M = marker).

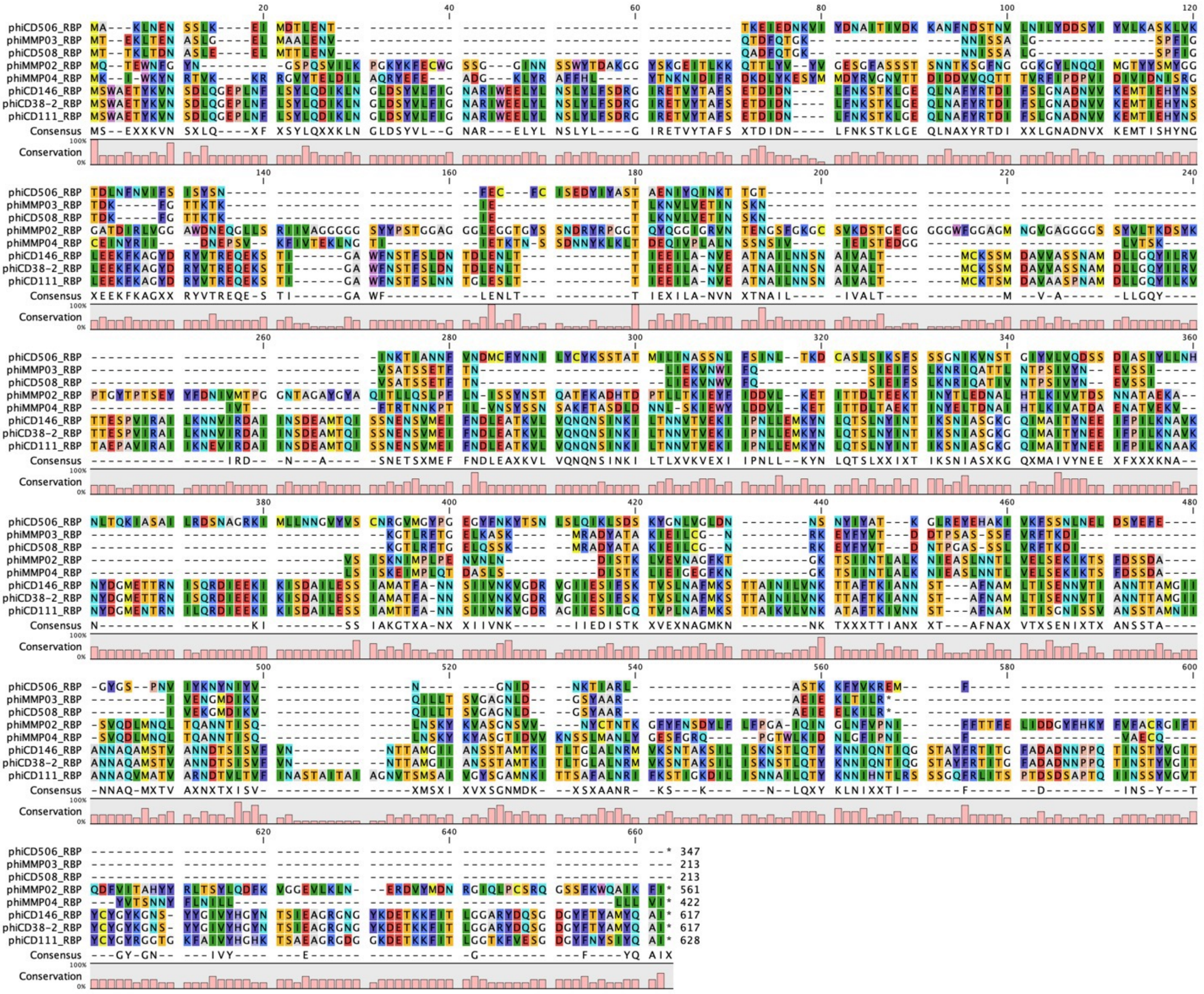


**Supplementary figure 3.** Multiple protein sequence alignment of the predicted phage RBPs.

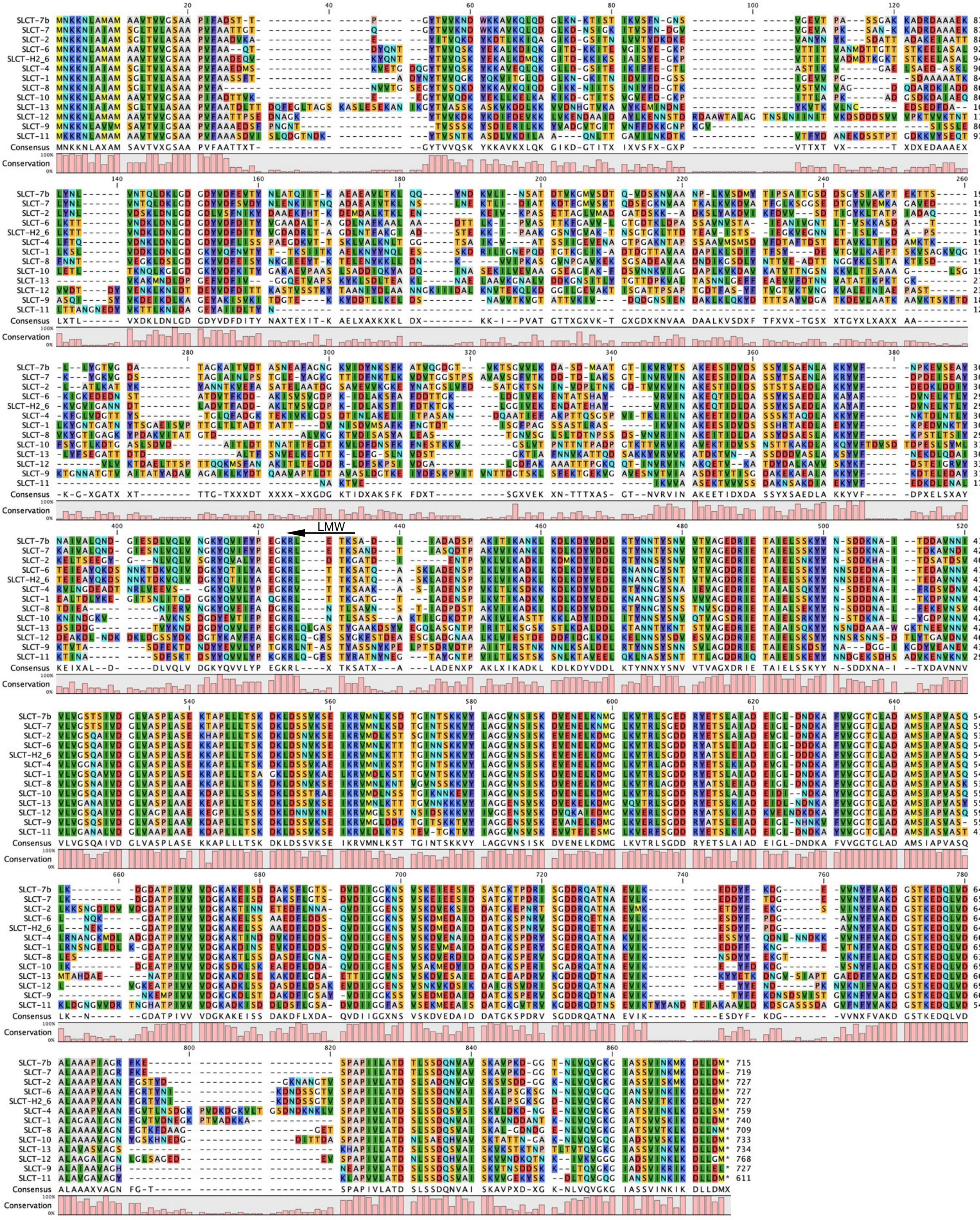


**Supplementary figure 4.** Multiple protein sequence alignment of the full-lengt SLCTs used in this study. The end of the N-terminal LMW fragment is indicated by an arrow (the exact position varies in function of the SLCT).

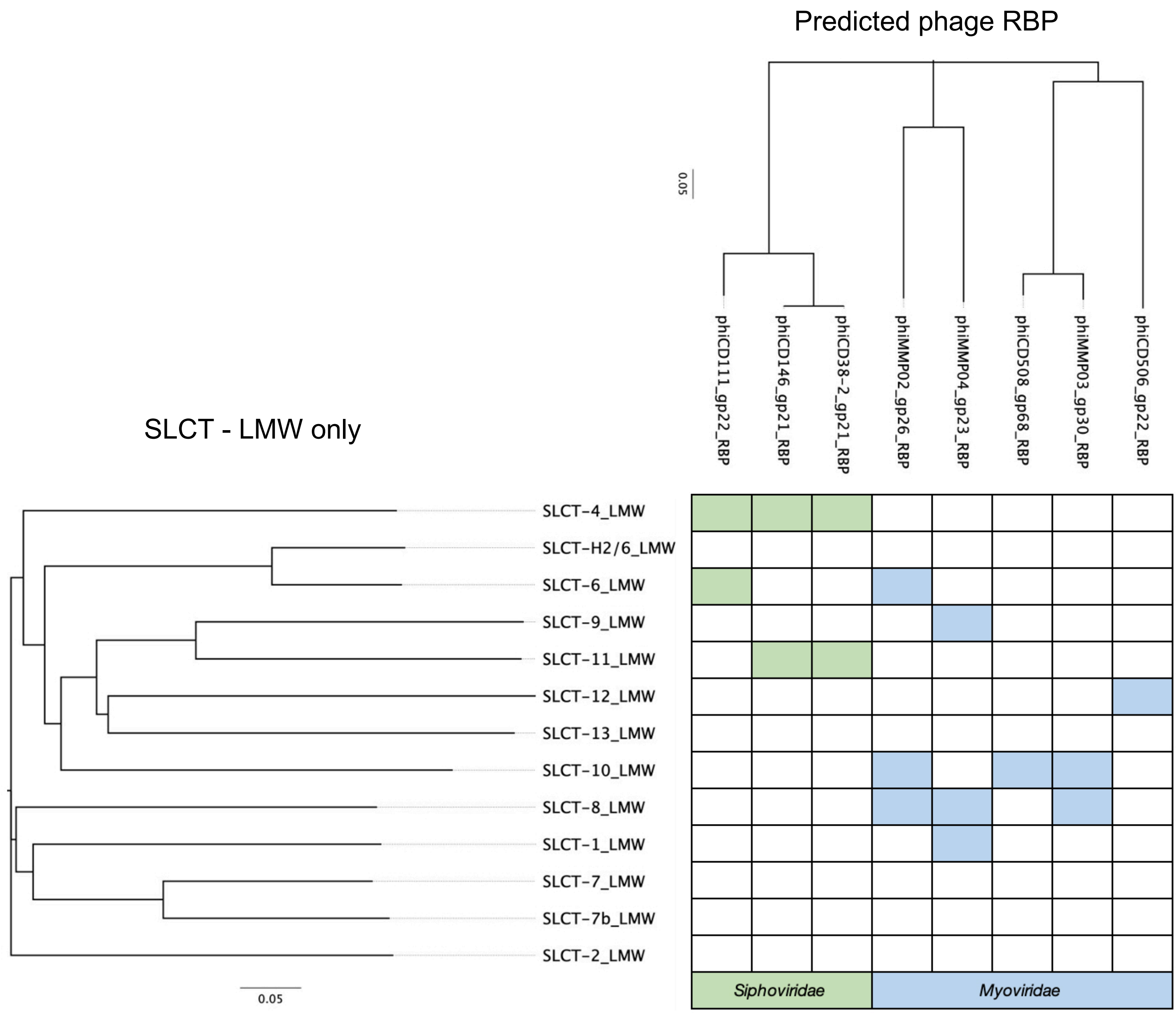


**Supplementary figure 5.** Relation between SLCT, phage RBP and host susceptibility to phage infection. Multiple alignments and phylogenetic trees of the LMW fragment from each SLCT and the predicted phage RBPs were compared with the host range of the phages (see Table 2 for details).
